# *Aeromonas hydrophila* induces skin disturbance through mucosal microbiota dysbiosis in *Pangasianodon hypophthalmus*

**DOI:** 10.1101/2022.01.27.478127

**Authors:** Li-Hsuan Chen, Chia-Hsuan Lin, Ru-Fang Siao, Liang-Chun Wang

## Abstract

Bacterial pathogens are well-equipped to adhere to and initiate infection in teleost fish. The fish skin mucus serves as the first barrier against environmental pathogens. The mucus harbors commensal microbes that form the microbiota, impacting the host physiological and immunological regulation. However, how the skin mucosal microbiota responds to the presence of pathogens remains largely unexplored. Thus, little is known about the status of skin mucus prior to the infection with noticeable symptoms. In this study, we aim to investigate the interaction between pathogen and skin mucosal microbiota, as well as the fish skin immune responses in the presence of pathogens. By challenging striped catfish (*Pangasianodon hypophthalmus*) with different concentrations of bacterial pathogen *Aeromonas hydrophila* (AH), the skin immune response and the mucosal microbiota were examined by qPCR and 16S rRNA gene sequencing analysis. We found a pathogen concentration able to stimulate the skin immune response associated with the significant mucosal microbiota change and re-confirmed with the *ex vivo* fish skin model. Further analysis indicated that the change was attributed to the significant increase in opportunistic pathogens over AH. We concluded that the presence and increase of AH results in dysbiosis of mucosal microbiota that can stimulate skin immune response. We believe our work can shed some light on host-pathogen-commensal microbiota interaction and therefore contribute to aquaculture infection prevention.

**IMPORTANCE:** The fish skin mucosal microbiota is essentially the first barrier in response to the presence of pathogens. This study is the first study to elucidate the interaction between the AH, the skin mucosal microbiota, and the striped catfish skin at the initiation stage of infection. Our study provides a platform to study both the correlation and causation of the interaction between pathogen, the fish skin, and the skin mucosal microbiota. This work provides information that changes in the AH-induced mucosal microbiota result in skin disturbance with immune stimulation.

## INTRODUCTION

Fish skin serves as the first defense line against pathogens that plays vital functions in the immunity, osmoregulation, and endocrine signaling of fish (1). The outer layer of the fish skin is the mucus layer secreted by the epidermal goblet cells (2). The mucus is vital for the immune system and protects against pathogens by physical, chemical, and biological implements (3–5). The physical and chemical properties allow mucus to continuously sloughed off and contain host-secreted antimicrobials such as lysozyme, immunoglobulin, and lectins (5, 6). However, the biological property also has an interactive and complex role in regulating homeostasis (7, 8). The microbiota, defined as the microbial community with distinct physio-chemical properties, has evolved to reside with the host and can regulate host-pathogen development. The regulation of the microbial community in mucosal epithelium includes enhancement of the epithelial barrier, development of the immune system, and nutrient acquisition (9–12). In aquaculture, although the role of mucosal microbiota has shown its importance against pathogens by the stimulation of the immune system (13, 14), the complex interaction of mucosal microbiota with the pathogens and fish skin immune system is still unexplored.

*Pangasianodon hypophthalmus*, commonly named striped catfish, is a major freshwater species and dominants 2% of global live-weight aquaculture production (15). Nonetheless, bacterial pathogens can cause production loss (16, 17). The bacteria that commonly cause catfish disease is *Aeromonas hydrophila* (AH) (17–20). AH is the causative agent of Motile Aeromonas Septicemia (MAS) in the farmed striped catfish, especially in the juvenile stage (21). Studies have reported the molecular responses of catfish after bacterial infection and described genomic and genetic bases responsible for disease development (22). Studies have also shown that catfish with mucus scraping could successfully initiate AH infection while an undisturbed mucus did not result in AH infection regardless of any dose or presence of AH (23, 24), hinting an essential role of skin mucus in interfering with pathogen entry in the natural state. However, the initial interaction between mucus and pathogen along with skin has not assessed. Therefore, this study aims to mimic the natural state to understand the events on the skin mucus prior to potential infection or damage.

## RESULTS

### The immune stimulation was found at skin tissue in response to AH challenge

The skin is the primary portal responding to the presence of AH. A study has reported multiple immune markers regulated in the skin during AH challenge in channel catfish (24). To validate the host immune response in the striped catfish skin under our AH challenge setting, Quantitative Real-Time PCR (qPCR) was used to measure the expression level of targeted immune genes in the skin (Fig. 1A). In general, we found a significant immune stimulated response only at 10^6^ AH/ml group compared to the non-AH control (Fig. 1B-H). The immune stimulated response in 10^6^ AH/ml group had a 4 to 8-fold increase in TLR4, TLR5, MyD88, IL-1β, IL-10, and Mucin-5AC expression compared to the non-AH control. Moreover, a 32-fold increase in IL-8 has been observed (Fig. 1F). Additionally, we found a gradual but not threshold-dependent increase in IL-10 and Mucin-5AC expression when comparing 10^6^ AH/ml to the rest of the groups (Fig. 1G&H). To further exclude the possibility that this immune stimulatory response was systemic, IL-1β and IL-10 expression levels were examined in the liver, spleen, and kidney from fish in 10^6^ AH/ml group and compared to those from the non-AH control. We found no significant difference in IL-1β and IL-10 expression levels between the two groups (Fig. S1), indicating the immune stimulation was not systemic. Additionally, all fish in the two groups survived without noticeable symptoms and the presence of AH in the liver, spleen, and kidney (Table S1). These data suggest that the skin with the presence of mucus can be immune stimulated by continuous AH bath challenge.

**FIG 1.**
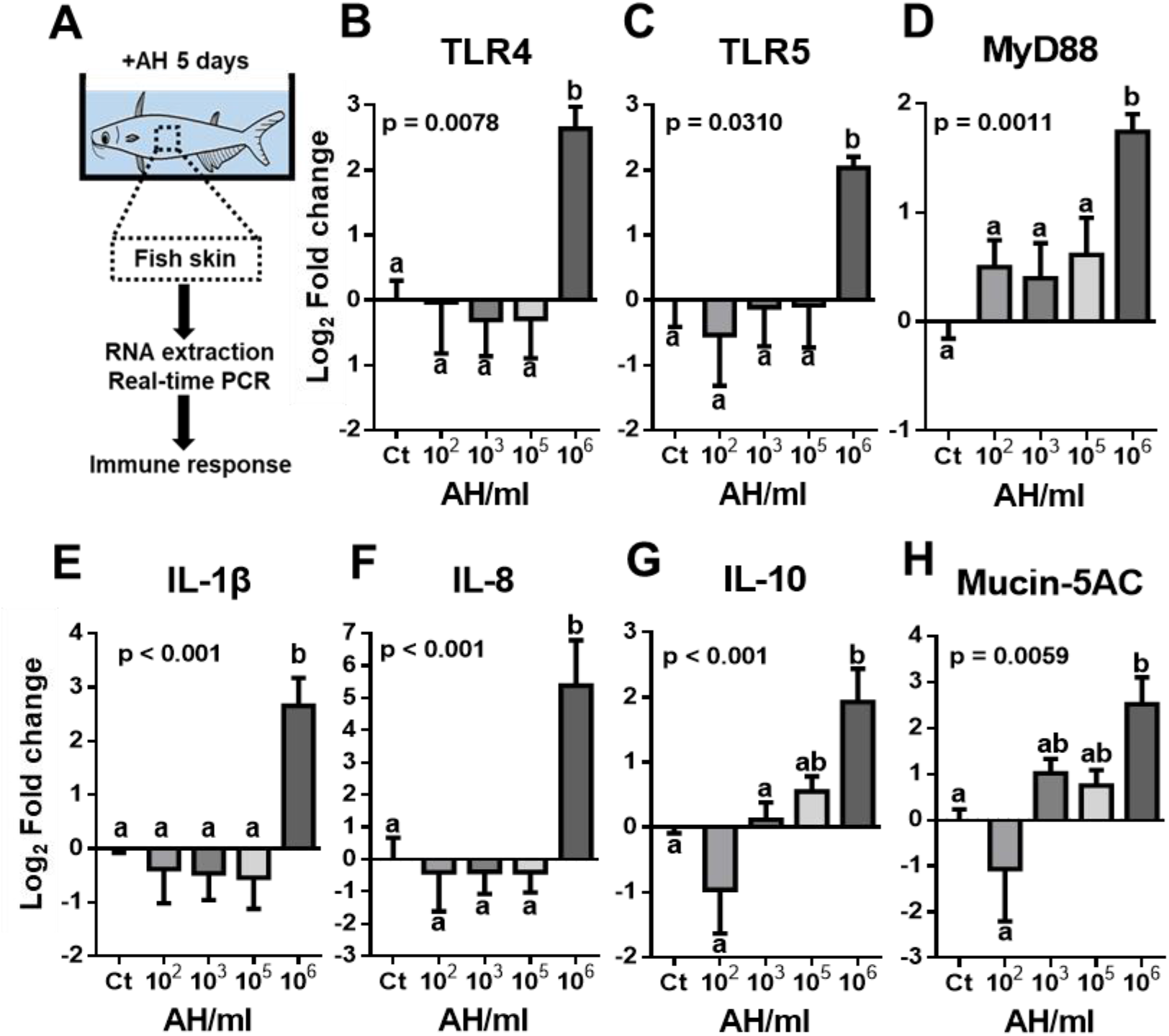
Fish skin immune response after AH challenge. qPCR of AH-challenged and non-challenged fish skin was performed to quantify the immune response. (A) Illustration of the experiment. Expression level of (B) TLR4, (C) TLR5, (D)MyD88, (E) IL-1β, (F) IL-8, (G) IL-10 and (H) Mucin-5AC were shown. N=8 fish in each group. Statistical significance was determined by One-way ANOVA followed by Tukey’s post-hoc test. (Different letters indicate significant differences at p < 0.05)

### AH-challenged skin mucus elicits similar immune stimulation in the *ex vivo* skin model

Studies have shown an essential role of fish skin mucus in stimulating and regulating the immune response (6, 25). Based on this, we hypothesized that the AH-challenged skin mucus could induce the observed skin immune response. To test this hypothesis, an *ex vivo* skin model (26) was used to create an interactive environment between the skin and the mucus (Fig. 2A). To first confirm the skin model can be used as a gnotobiotic tissue model, PCR of bacterial 16S rRNA gene V3-V4 region was performed on the culturing media, and the skin explant in the skin model. We found no bacteria in neither media from the apical and basal side of the model, nor the explant compared to a positive 500 bp band shown in fresh skin mucus (Fig. 2B), suggesting the skin model system can be used as a gnotobiotic tissue model. We then sought to examine the effects of mucus on the skin. Freshly collected skin mucus from AH-challenged and non-AH control fish were applied to the skin model system for 6 hr, and subsequent qPCR was performed to determine the expression level of IL-1β. We found that the expression level of IL-1β was an 8-fold increase in the skin model transplanted with mucus from 10^6^ AH/ml group compared to the non-AH control, 10^3^, and 10^5^ AH/ml groups, whereas no significant change was found among the other comparisons (Fig. 2C). To further examine whether the increased expression of IL-1β resulted from living biological properties, a frozen mucus of the 10^6^ AH/ml group was examined. Without any growth of bacteria from frozen mucus (data not shown), we found no significant IL-1β expression change in the skin applied by the frozen mucus compared to fresh mucus (Fig. 2D), indicating the living biological property can cause the immune stimulation. These data suggest that the living biological property of AH-challenged skin mucus can directly stimulate the skin immune response, similar to the response observed *in vivo*. Therefore, the mucus with its living biological property under the interaction with AH may play a vital role in the health status of the fish skin.

**FIG 2.**
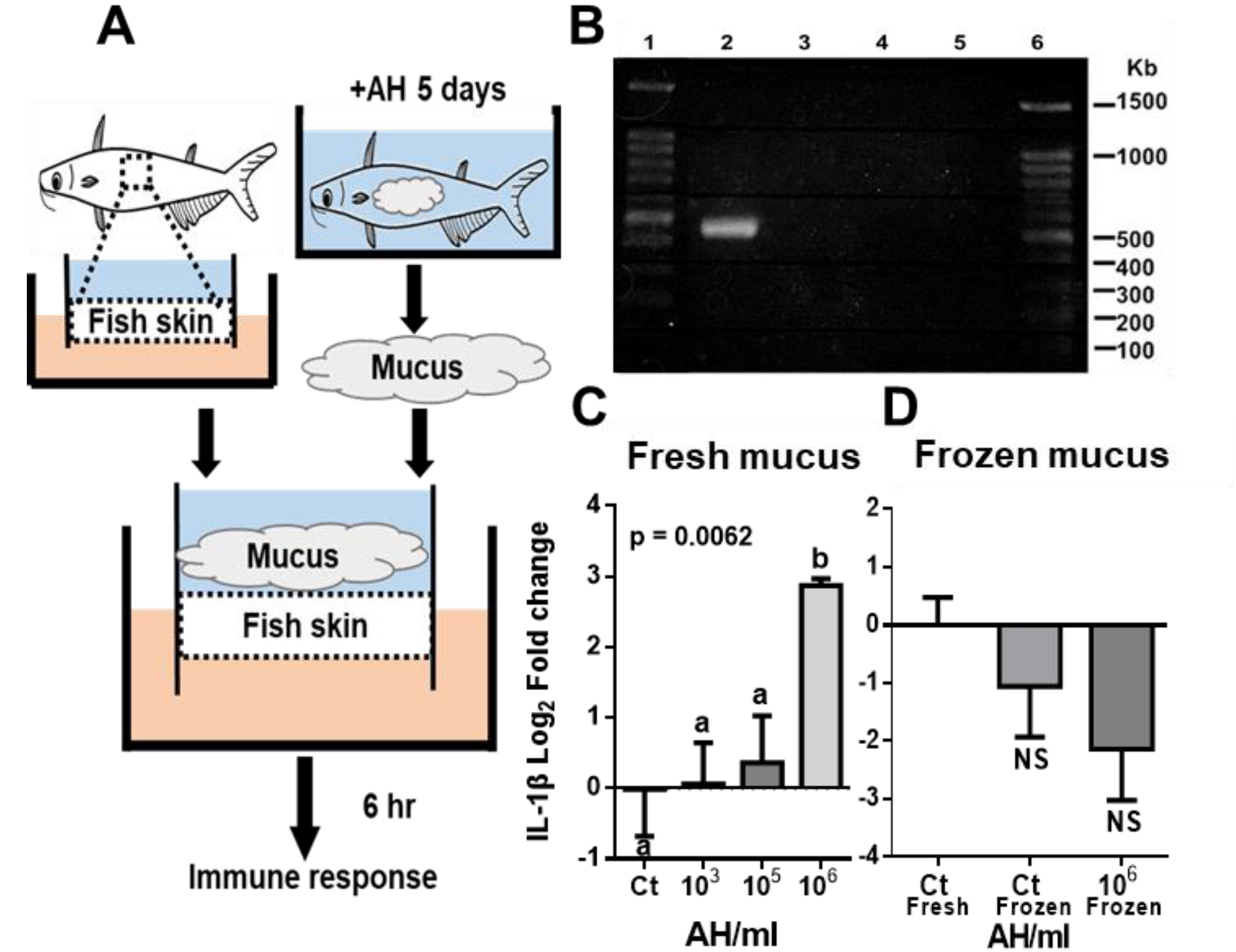
Fish skin immune response of the *ex vivo* skin model inoculated with mucus from AH-challenged fish. The skin mucus of AH-challenged and non-challenged fish was scraped and transplanted to the *ex vivo* skin model for 6 hr. qPCR of the skin tissue was performed on IL-1β expression level. (A) Illustration of the experiment. (B) Confirmation of the gnotobiotic property in skin model by gel electrophoresis of amplified 16S rRNA gene V3-V4 region. (Lane 1 and 6 : 10 kb ladder; Lane 2 : Positive control (AH); Lane 3: Negative control (double-distilled water); Lane 4:media from the apical and basal side of the model; Lane 5:Skin from the model system). (C) IL-1β expression level of the *ex vivo* skin with (C) fresh mucus and (D) frozen mucus from AH-challenged fish. N=5 fish in each group. Statistical significance was determined by One-way ANOVA followed by Tukey’s post-hoc test. (Different letters indicate significant differences at p < 0.05)

### The skin mucus mixed with AH does not stimulate the skin immune response

The microorganism is the critical living biological property in the skin mucus. Microbial factors, such as pathogens and their toxins, in the mucus have shown to perturb the host immune system (27). The presence of AH has shown to stimulate the skin immune response. However, whether AH in the mucus can stimulate the skin immune response is unknown. We addressed this question by inoculating different numbers of AH into the mucus collected from non-challenged fish skin and applying them to the skin model. To mimic condition *in vivo*, the viable AH concentration in the mucus was determined by diluting and plating the freshly collected mucus from AH-challenged and non-AH control fish on Starch ampicillin agar (Fig. 3A). The counting method was performed by adding Iodine solution to distinguish AH from other bacteria by starch hydrolysis around the AH colonies (Fig. 3B). We found viable AH in every challenged group but not in the non-AH control. In the challenged groups, the AH concentration in the mucus was increased as the challenged concentration increased but was 5 to 10-fold lower to the challenged concentration in the water tank. In addition, similar AH concentrations in the mucus can be found at 10^2,^ and 10^3^ AH/ml challenged concentrations (Fig. 3C).

**FIG 3.**
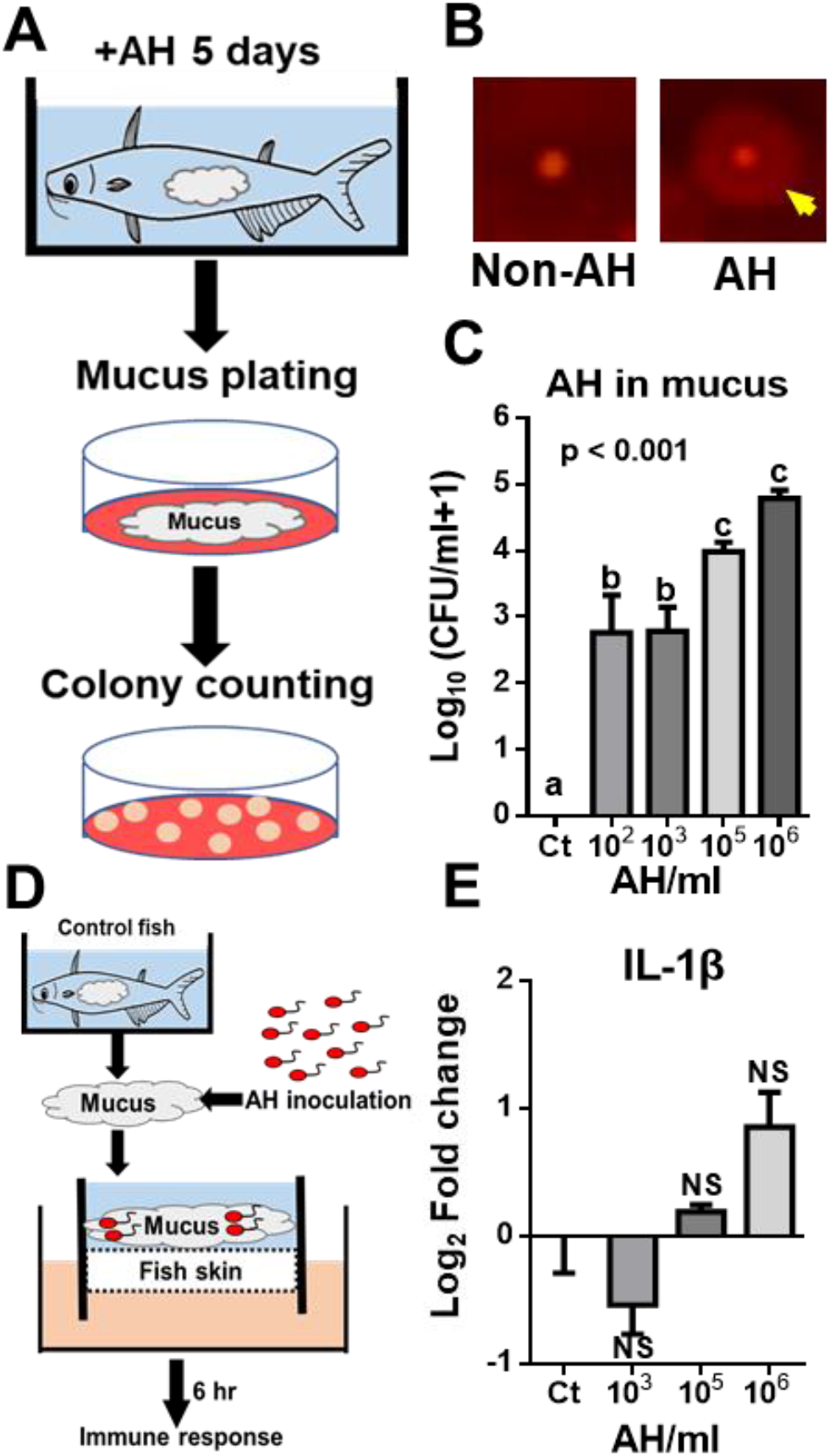
Viable AH in mucus and its contribution to fish skin immune response in the *ex vivo* skin model. Viable AH concentration in mucus was quantified by directly plating collected skin mucus. (A) Illustration of viable AH quantification. (B) Starch hydrolysis of AH and non-AH bacteria in Starch Ampicillin Agar with iodine solution. Yellow arrow indicates the clear halo surrounding the colony due to starch hydrolysis. (C) AH survival in mucus was shown as colony-forming unit (CFU) per milliliter mucus. N=8 fish in each group. Comparable number of AH was mixed into the mucus from non-challenged fish and inoculated on the skin of the *ex vivo* skin model for 6 hr. qPCR was performed for IL-1β expression level. (D) Illustration of the AH-mucus mix in the *ex vivo* skin model. (E) IL-1β expression level of the *ex vivo* skin with AH-mucus mix after 6 hr incubation. N=5 fish in each group. Statistical significance was determined by One-way ANOVA followed by Tukey’s post-hoc test. (Different letters indicate significant differences at p < 0.05)

To then examine whether the corresponding concentration of AH in the mucus can induce the skin immune response, the desired AH number was inoculated into the collected skin mucus from non-challenged fish to create the AH-mucus mix without changing the rest microbial community. The mucus was then applied to the skin model and incubated for 6 hr followed by measuring the IL-1β expression level (Fig. 3D). We found no significant change of IL-1β in the skin model applied with the AH-mucus mix corresponding to concentration *in vivo* compared to the skin model with control mucus. (Fig. 3E). These data suggest that the concentration of AH in the mucus under our challenge range alone may not contribute to the host skin response. Therefore, it implies that the interactive biological property, instead of the AH itself, potentially contributes to the skin immune response.

### The compositional change of the skin mucosal microbiota by AH challenge

The microbiota is the microbial community playing a fundamental role in maintaining the health of teleost fish, mediating the interaction between the pathogen and the host immune system (13, 28). Since the immunostimulatory effect was not elicited by mucus mixed with AH alone but the mucus from AH-challenged fish, we hypothesized that the skin mucosal microbiota underwent the significant change by AH challenge. To examine if the skin mucosal microbiota composition changed by AH challenge, the 16S rRNA gene V3-V4 region of the collected skin mucus of individual fish from AH-challenged and non-AH control were sequenced for determining the bacteria taxonomy, richness, and abundance (Fig. 4A). In Alpha-diversity, we found the 10^6^ AH/ml, but not other AH-challenged groups, had a significantly higher Chao 1 index than the control (Fig. 4B). However, we found a higher but not significant level in the Shannon index between each group (Fig. 4C). In Beta-diversity, Principal Coordinates Analysis (PCoA) analysis was performed to examine the similarity among individuals and groups. In general, we found that individuals in the 10^2^ and 10^3^ AH/ml challenged group clustered together with non-AH control (Left-hand side of Fig. 4D) whereas individuals in the 10^5^ and 10^6^ AH/ml AH-challenged group clustered together but away from the non-AH control (Right-hand side of Fig. 4D). To further confirm the clustering difference in PCoA, Bray–Curtis dissimilarity boxplot was generated and confirmed the significant difference between the clustering of the non-AH control, 10^2^, and 10^3^ AH/ml challenged group, and the clustering of the 10^5^ and 10^6^ AH/ml challenged group (Fig. 4E). These data suggest the microbiota composition changed significantly in the 10^5^ and 10^6^ AH/ml AH-challenged group. We denote 10^5^ and 10^6^ AH/ml as high concentration groups, whereas 10^2^ and 10^3^ AH/ml as low concentration groups. To examine the compositional difference of microbiota between the two clustering groups and compared to the non-AH control, microbiota composition and abundance were classified into phylum and family level for comparisons. At the phylum level, we found minimal difference where the most abundant phylum in all samples were Proteobacteria, Firmicutes, and Actinobacteria, comprising more than 95% of the microbiota (Fig. 4F & S2A). However, a drastic difference was observed at the family level in high concentration groups compared to other groups (Fig. 4G & S2B). The most abundant bacteria family at non-AH control and low concentration groups were similar, and Burkholderiaceae, Paenibacillaceae, and Xanthomonadaceae occupied 60-70% abundance. However, in high concentration groups, we found 40-50% lower in Burkholderiaceae while 12-28% higher in Xanthobacteraceae followed by a 5-10% increase in Vibrionaceae, and 2-5% increase of Enterobacteriaceae, Corynebacteriaceae, Moraxellaceae, and Caulobacteraceae. Therefore, the AH challenge induced a significant change of skin mucosal microbiota at a high challenging concentration.

**FIG 4.**
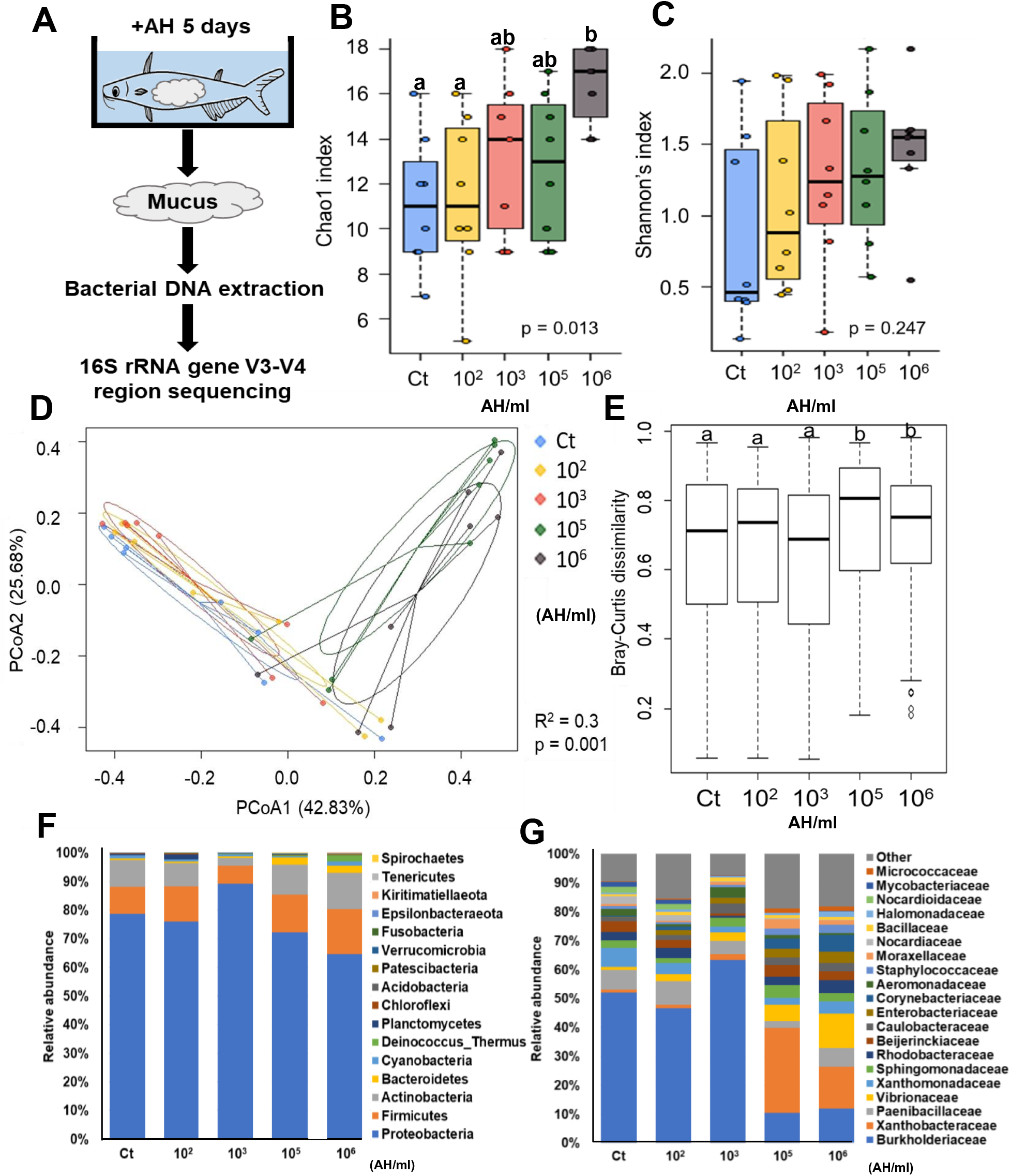
Changes in the skin mucosal microbiota after AH challenge. The skin mucus of AH-challenged and non-challenged fish was scraped, and the microbial DNA was extracted, followed by 16S rRNA gene sequencing. (A) Illustration of the experiment. The alpha-diversity indices:(B) Chao 1 and (C) Shannon. (D) Principal coordinate analysis (PCoA) and (E) Bray–Curtis dissimilarity index was generated to determine the beta-diversity at family level. N=8 fish in each group. The relative abundance of the skin mucosal microbiota composition was shown at (F) phylum level and (G) family level. Statistical significance was determined by One-way ANOVA followed by Tukey’s post-hoc test. (Different letters indicate significant differences at p < 0.05)

### Microbial composition change at high challenge concentration alters the functional capacity and metabolic output

When microbial composition changes, the metabolic output and functional capacity of the community may also alter (29). To determine if the metabolic output and/or function capacity changed in skin mucus microbiota, Phylogenetic Investigation of Communities by Reconstruction of Unobserved States (PICRUSt2) analysis using Kyoto Encyclopedia of Genes and Genomes (KEGG) database was performed to compare microbiota metabolic output and function from different AH-challenge groups to the non-AH control. The significant difference and abundance of KEGG pathways analyzed by One-way ANOVA are summarized in Fig. 5. The level 1 pathways (Fig. 5A) indicated a general metabolic output such as the metabolism of terpenoids, polyketides, xenobiotics, lipid, amino acid, and carbohydrate, the biosynthesis of other secondary metabolites, nucleotide replication, repair, and membrane transport. The significant difference in relative abundance of sub-categorized level 2 pathways (Fig. 5B & 5C) was specifically analyzed between high concentration groups and the rest of the groups (Tukey’s post-hoc test, p<0.05). We found a significant increase of relative abundance related to linoleic acid, cyanoamino acid metabolism and nitrotoluene degradation while a significant decrease of relative abundance related to bacterial secretion system and ascorbate and aldarate metabolism, suggesting that the functional capacity and metabolic output of the skin mucus microbiota were altered by AH challenge.

**FIG 5.**
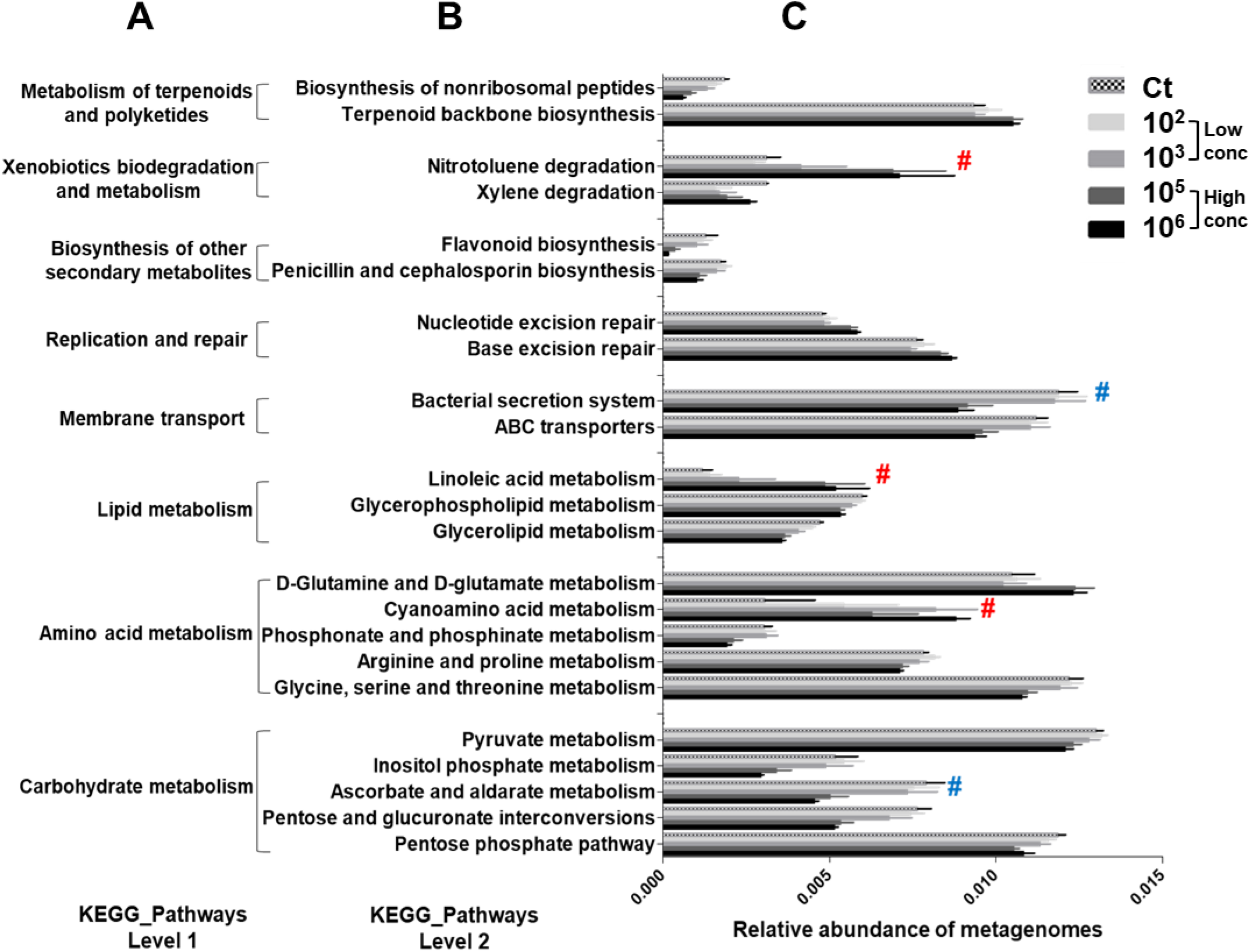
Changes in the metabolic and function output of skin mucosal microbiota after AH challenge. Functional prediction of the microbiota was performed by PICRUSt2 and annotated by the KEGG database. The significant difference in (A) Level 1 and (B) Level 2 of KEGG pathway was shown. (C) The relative abundance of each functional pathway. N=8 fish in each group. Statistical significance was determined by One-way ANOVA followed by Tukey’s post-hoc test. (# indicates significant differences at p < 0.05 in the high concentration group compared to the low concentration group and non-AH control. Red # indicates up-regulated pathway while blue # indicates down-regulated pathway.)

### The rise of opportunistic pathogens in mucosal microbiota potentially stimulates the skin immune response

The composition and function analysis have shown the significant difference in skin mucosal microbiota between low and high AH concentration challenged groups. However, the immune markers expression level significantly increased only at 10^6^ AH/ml in high AH concentration challenge groups *in vivo* and *ex vivo*. We then hypothesized a difference in skin mucosal microbiota between 10^5^ and 10^6^ AH/ml challenged group in determining the stimulation of the skin immune response. Since the two groups were not significantly different in Bray-curtis matrix, we specified the microbiota difference by performing the Linear discriminant analysis Effect Size (LEfSe) analysis to show the unique niche of bacteria group at 10^6^ AH/ml challenged skin mucosal microbiota compared to the rest of the groups. We found a unique niche of *Vibrio, Corynebacterium, Paracoccus, Brevundimonas,* and *Escherichia_Shigella* at 10^6^ AH/ml (Fig. 6A). We then correlated these unique bacteria genera to the increase of IL-1β expression level and found a significantly positive correlation (r=0.5, p=0.0011) (Fig. 6B) Moreover, we found the relative abundance of these genera was gradually increased when challenge concentration increased and peaked at 10^6^ AH/ml (Fig. 6C). Among these genera, *Vibrio* has the highest increase from 1% in non-AH control to 5% in 10^5^ and 10% in 10^6^ AH/ml group. Other genera, such as *Corynebacterium_1, Paracoccus, Brevundimonas,* and *Escherichia_Shigella* have a gradual with 2 to 4% relative abundance increase through low to high AH challenge group. Additionally, we found only *Vibrio,* but no other genera have a significant increase of relative abundance in 10^6^ AH/ml group compared to non-AH control while there is no significant difference between 10^5^ AH/ml group and non-AH control in these genera by Student’s t-test in Table S2. We then hypothesized 10^6^ AH/ml challenge concentration can induce the growth of genus *Vibrio* over a threshold concentration to induce the IL-1β expression level. We compared the concentration of AH and *Vibrio* in the skin mucus of each fish in different groups using the viable AH and the abundance from plating and the 16S rRNA analysis. We found *Vibrio* had a significant 10-fold higher than AH in the 10^6^ AH/ml group, whereas a non-significant difference was observed in other groups (Fig. 6D). The data suggest that *Vibrio* can increase to the number over AH in the mucus and potentially be the stimulant of skin immune response. Taken together, the increase in AH concentration can concomitantly increase the opportunistic pathogens, which may play a primary role over AH in stimulating the host immune response.

**FIG 6.**
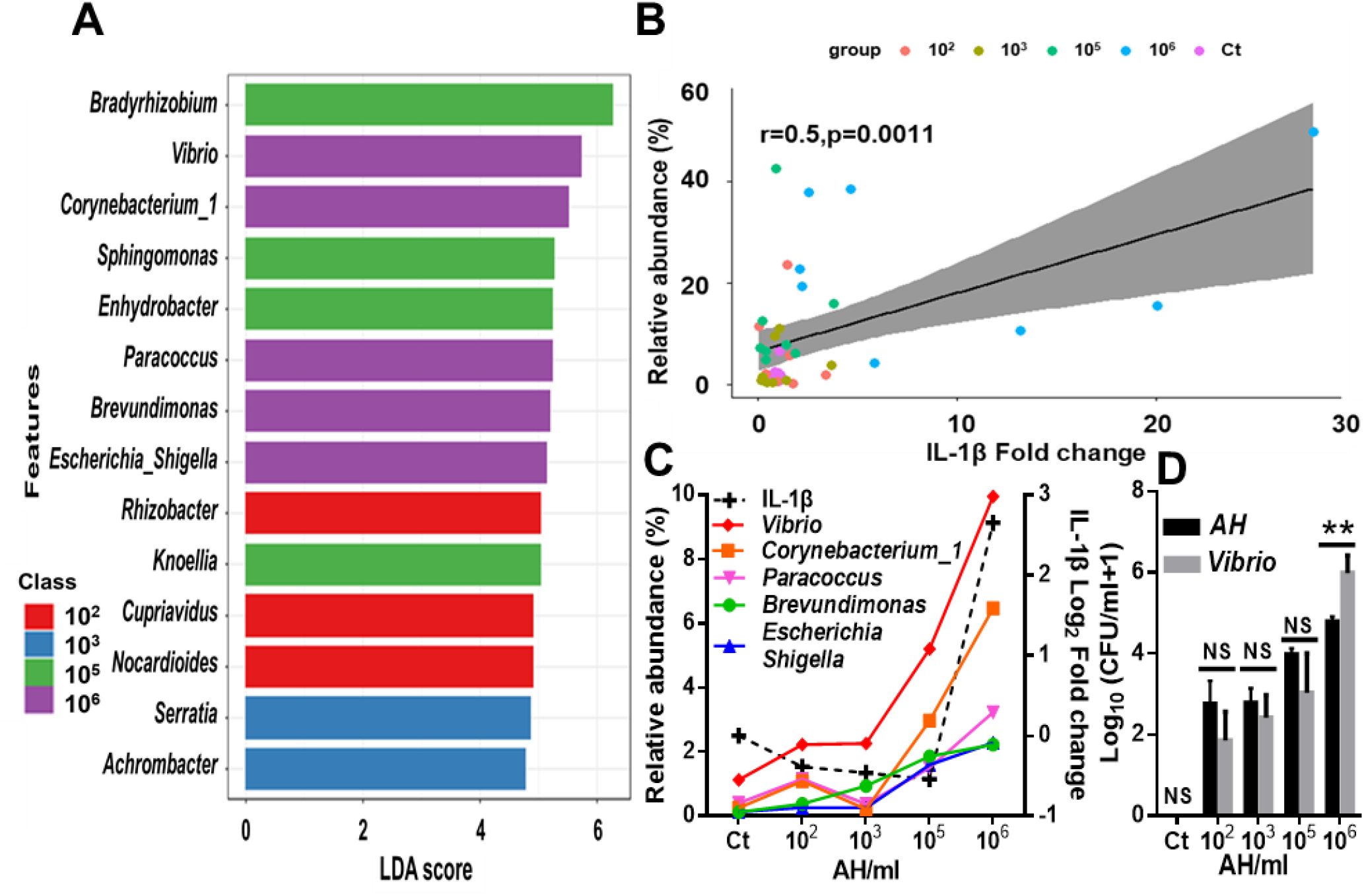
An increase of opportunistic pathogens correlates to skin immune stimulation. Comparison of the skin mucosal microbiota composition of 10^6^ AH/ml group to the rest of the groups. (A) LEfSe analysis shows differentially abundant genera as biomarkers determined using Kruskal-Wallis test (p-value cutoff = 0.1) with LDA score = 2. (B) Correlation between IL-1β expression level and the increased skin mucosal microbiota genera in 10^6^ AH/ml-challenged fish. (C) Relative abundance trend of increased genera in 10^6^ AH/ml group followed by increased AH-challenged concentration (D) Comparison of viable *Aeromonas hydrophila* and *Vibrio* in mucus under different AH-challenged concentrations. N=8 fish in each group. Statistical significance was determined by Student’s t-test. (***p ≤0.001; **p≤0.01; *p≤0.05)

## DISCUSSION

Initial interaction between pathogen, skin mucosal microbiota, and the host skin is essential but largely unexplored. In this study, we showed the presence and increase of AH results in dysbiotic mucosal microbiota that can stimulate immune response of the skin (Fig. 7).

**FIG 7.**
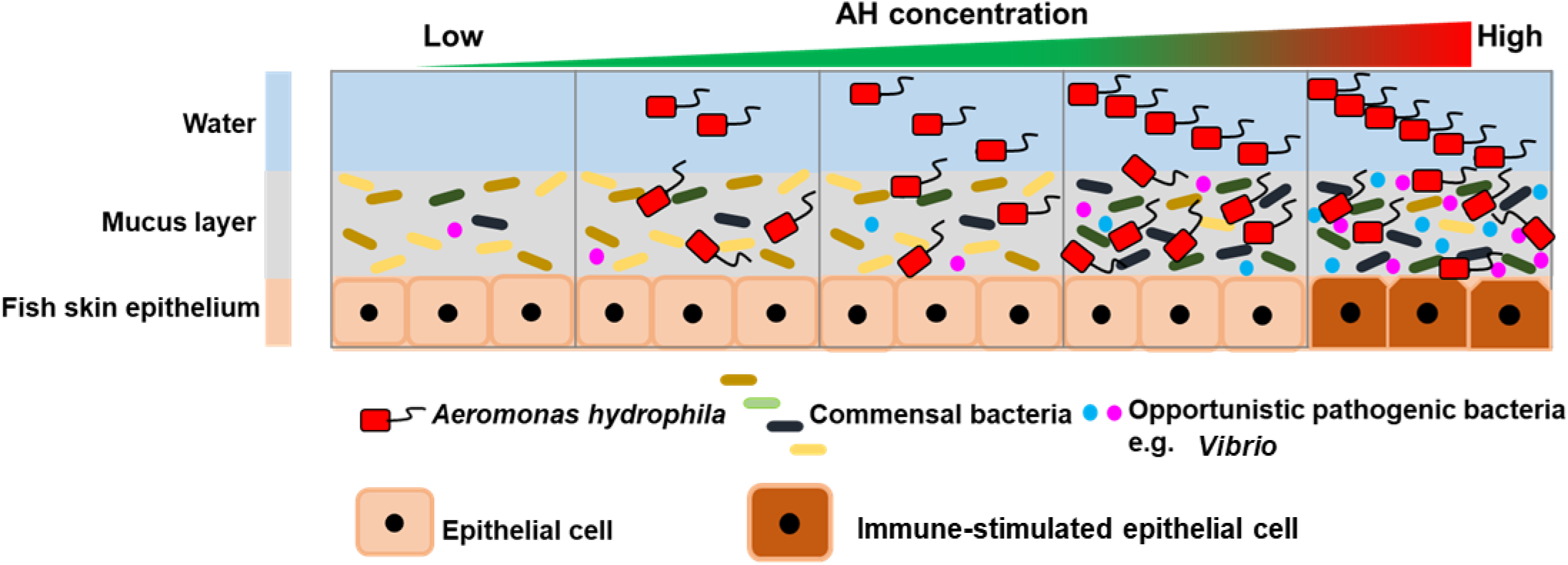
A prospective model of interactions between host skin, mucosal microbiota, and AH. The results suggest that the fish skin mucosal microbiota composition is maintained and presents homeostasis in the mucosal layer in the presence of a low concentration of AH. With an increased concentration of AH, the skin mucosal microbiota composition changes but still maintains homeostasis in the mucosal layer. Once the AH concentration increases over a critical threshold where opportunistic pathogens rise in the changed microbiota, dysbiosis occurs in the mucosal layer thus stimulating immune response in the skin.

Studies have shown that fish skin mucus is essential for regulating skin immunity and protecting fish from pathogen invasion (30). In our study, performing *in vivo* experiments and using an *ex vivo* skin model, we found the stimulated immune response in the skin model inoculated with the fresh challenged mucus (Fig. 2C) but not the un-challenged mucus mixed with AH or the frozen mucus (Fig. 2D&3E). Our previous study observed a significant immunostimulatory effect when inoculating the same amount of AH directly on the skin model without mucus (26). Therefore, the striped catfish skin mucus has a protective role in response to AH. A similar observation has been claimed in blue and channel catfish studies, where infection occurred when the skin mucus was abraded/scraped off under bath challenge (23, 24). Several studies suggested that the skin mucus possesses host-secreted antimicrobial properties that protect the host from pathogens (31, 32). We have explored the anti-AH property of the skin mucus in striped catfish but found no effect against AH growth (Table S3). A similar observation has been found using the apical media containing the mucus secretion from the *ex vivo* model (data not shown). This suggests the skin mucus possesses an immune-buffering rather than growth-inhibitory role against AH, potentially from the microbiota. Two possible mechanisms can be hypothesized. First, the skin mucosal microbiota can direct buffer the immune response by interacting with AH. Several Bacteria can serve as potential candidates in this buffering system by colony resistance and/or production of adherence-inhibitory compounds (33–35). Second, previous studies have shown that fish commensal microbiota can regulate the host immune system (28, 36–39). An immunostimulatory role was also found within the striped catfish skin mucus using the *ex vivo* skin model (Fig. S3). Multiple bacteria genera can stimulate and boost the host immune system for responding to pathogens faster. These bacteria genera include *Lactobacillus*, *Shewanella*, and *Bacillus* (40–42). Even though these bacteria genera are not in the feature table of our research, changing the property of mucosal microbiota may be enough to induce immune stimulation (9). It would be interesting to manipulate the mucosal microbiota composition using mucus isolates and metabolism under different stress conditions to examine how the mucosal microbiota can change the skin immune response.

From the *in vivo* and *ex vivo* model, we have gained mechanistic information on how AH challenge induces the rise of pathogens in mucosal microbiota that can stimulate immune response of the skin (Fig. 7). Here, we found *Vibrio,* which has been reported as an important opportunistic pathogen responsible for aquatic infectious diseases (43–45), drastically increases in the skin mucus of 10^6^ AH/ml challenged fish. This suggests AH can directly or indirectly trigger the growth of the opportunistic pathogen. Rolig et al. has shown that the presence of AH can increase the number of *Vibrio* and attract neutrophils in the gut of gnotobiotic zebrafish (38). This data supports a hypothesis that AH can directly help the increase of *Vibrio*. Zhang et al. has also shown that the presence of parasitic pathogen infection can influence the skin microbiota and change the skin mucus into an opportunistic pathogen favored environment in rainbow trout (46). However, the other hypothesis on the host contribution to the rise of *Vibrio* is also possible. Studies have shown that the teleost stress itself can influence the skin microbiome and change the skin mucus into an opportunistic pathogen favored environment in Brook trout (47). Both mechanisms can occur individually or synergistically on the fish skin. In addition to *Vibrio*, we have also noticed other genera increased at 10^6^ AH/ml compared to 10^5^ AH/ml. *Corynebacterium_1, Paracoccus, Brevundimonas, and Escherichia_Shigella*, were present in the low concentration challenge group or non-AH control and then gradually increased when challenge concentration increased. Studies have shown that these bacteria can be opportunistic aquatic pathogens. *Corynebacterium_1* was isolated from the diseased striped bass (48), *Escherichia_Shigella* is a common opportunistic pathogen in the gastrointestinal tract of virus-infected fish (49), and *Brevundimonas* has recently been defined as an emerging global opportunistic pathogen (50). Therefore, it is reasonable to postulate that the mucus changed by AH challenge might become an environment for these opportunistic pathogenic bacteria genera to become noticeable in abundance. In addition, the PICRUSt2 analysis of metabolic profile also showed the mucus environment from high concentration challenged groups favors microbiota with nucleotide repair and linoleic acid metabolism (Fig. 5), implying an unfavorable environment for microbial growth (51, 52). With *Vibrio* and other halophiles growing, a hypothesized higher osmotic environment can be in the skin mucus in the high concentration challenged fish. Studies have found that the presence of pathogens or other stress can regulate fish skin osmolality, driving salt secretion (53). It would be interesting to examine Na^+^/K^+^-ATPase activity and salt density in mucus to correlate the extremophiles we found. Taken together, the tripartite interaction of the pathogen, the skin, and the skin mucosal microbiota is essential and inseparable when interpreting the infection and inflammation.

Despite the broad information, this work provides to this tripartite interaction prior to or at the initial stage of AH infection, there are still several limitations in this work. First, the aquaculture environmental and seasonal factors can constantly change the skin mucosal microbiota (54), meaning the microbiota in the experimental fish in this study may only represent a niche of many cases. Previous studies have shown that fish skin mucosal microbiota represents major phyla, including Proteobacteria, Firmicutes, Actinobacteria, Bacteriodetes, and Cyanobacteria (55, 56). Our work has shown the same trend in phyla (Fig. 4F). However, this trend can be different in the family and genus level. Thus, our work may only apply to one of the several scenarios. Second, we found *Vibrio* contributes to the skin immune stimulation. Various species have been found as pathogens. *Vibrio* species, including *Vibrio harveyi, Vibrio owensii,* and *Vibrio parahaemolyticus* are major aquatic pathogens and were identified in our 16S analysis at species level (data not shown). However, 16S rRNA V3-V4 gene sequencing could not get accurate species-level identification. Isolation of targeted *Vibrio* and other potential opportunistic pathogens with further examination in the skin model with or without AH can be done in the future. Third, based on our data of increased IL-8, the neutrophil may be recruited. The previous study has shown that the immune cells work coordinately with epithelial cell signals to clear the pathogens (57). In the future, how the immune cells in the skin work can be explored by adding immune cells to our skin model. Fourth, the immune expression is the sum of the balance between immune-stimulation and –inhibition. Even though an immune stimulation was found only in 10^6^ AH/ml group after the five-day experiment, the regulation of immune system during the experiment and in the other groups were still unknown. The fluctuating expression of the innate immune markers in blue catfish was found in the first 12-24 hr after AH challenge, showing the fast immune response towards the presence of AH (24). It would be interesting to examine the immune regulation in a time-dependent manner during the challenge in the future.

In the future, we will explore the mutual interaction between each participant in this complicated interaction. On the bacteria side, the immunostimulatory roles of mucosal commensals and the mechanistic information on how AH induces the rise of opportunistic pathogens are of interest. On the host side, how the immune system in coordinated with the epithelial cells and mucus-secreting cells in response to the mucosal microbiota change induced by pathogens will need to be further understood. We believe this future work can shed some light on preventing early aquaculture infection.

## MATERIALS AND METHODS

### Experimental fish and model overview

*Pangasianodon hypophthalmus* were obtained from a local aquarium vendor and were bred and maintained in a 300 L tank at 28°C in the aquaculture room at the Department of Marine Biotechnology and Resources. A 12:12 hr light: dark period was maintained, and an air stone supplied supplemental aeration. After breading, eight fish with an average weight of 23 g were transited to separated 80 L tanks (40 L water) by experimental treatments and acclimated for half a month. After acclimation, fish in each tank were challenged with different pathogen *Aeromonas hydrophila* (AH) concentrations for 5 days. An overview of experimental treatments and transitions are given in Fig. S4. The water temperature during the challenge was controlled at 28°C. The experiment is permitted by the Institutional Animal Care and Use Committee of National Sun Yet-Sen University with approval numbers IACUC 10834 and 10836.

### Aeromonas hydrophila challenge

AH were cultured from a single isolate (AH20) (BCRC16704, Hsinchu, Taiwan) and were inoculated into SA agar (starch and ampicillin agar) (HIMEDIA M1177, India) and incubated in an incubator at 28°C overnight then bacterial cells were added to water to give a final concentration of 10^2^,10^3^,10^5^,10^6^ CFU/ml. The concentration of AH in each tank was determined using colony-forming unit (CFU) per ml by plating 100 ml of 10-fold serial dilutions onto SA agar plates every day after 1 and 24 hr after inoculation. To maintain the desired concentration in a five-day challenge period, tank water was changed and reinoculated with AH every day (Fig. S5 A-E). At day5 sampling day, the tank water microbial composition in the sampling day was analyzed to confirm the relative abundance of AH in each experimental treatment (Fig. S5 F-G).

### Skin tissue and mucus collection

Fish were collected from each of the appropriate control and treatment aquaria after a 5-day AH challenge. Fish were anesthetized by rapid chilling followed by cervical transection. The sampling procedures were conducted according to AVMA Guidelines for the Euthanasia of Animals. Skin mucus was removed by gentle scraping by slide. The fish skin tissue followed by liver, spleen, and kidney were then removed and flash-frozen in liquid nitrogen during collection and stored at -80°C until RNA extraction.

### Quantification of viable AH in mucus

The skin mucus was scraped after AH challenge. 20 μl mucus was mixed with 180 μl followed by serial 10-fold dilutions and plated onto SA agar plates. After incubated at 28°C overnight, the colonies were counted for calculating the actual concentration of AH in mucus. Iodine solution was added to the plate to distinguish AH from other bacteria by starch hydrolysis.

### Mucus transplantation

A striped catfish model system based on (26) was applied in this research. The fish skin tissue was first removed from the fish for making the *ex vivo* skin model and cultured in a CO_2_-free incubator at 25°C for 5 days. The detailed protocol is shown in Text S1. 20 μl of the skin mucus was scraped from challenge experiment or 20 μl of regular skin mucus mixed with desired AH in 2 μl were transplanted into the upper side of the fish skin model system with the double-distilled water on the apical side and the medium on the basal side. After 6 hr of mucus transplantation, the fish skin tissue was collected, followed by RNA extraction. cDNA library was built and applied to qPCR to quantify the desired immune markers.

### RNA extraction and Real-time PCR analyses

Each total RNA sample from the skin, liver, spleen and kidney of striped catfish were extracted with the TriPure Isolation Reagent (Roche, Mannheim, Germany) following the referenced instructions (58). cDNA library was built by M-MLV Reverse transcriptase (Promega, USA) following the manufacturer’s instructions. All the cDNA products were utilized for the quantitative real-time PCR reaction using the GoTaq qPCR Master Mix on a CFX96 real-time PCR Detection System (Bio-Rad, CA). Primer pairs were designed using Universal Probe library website and shown in Table S4. The comparative Ct (ΔΔCt) method was used to evaluate the expression of candidate genes (59). The detailed protocol is shown in Text S1.

### Skin mucus microbial DNA extraction, 16S rRNA gene sequencing and community analyses

Skin mucus microbial DNA was extracted. After preparation and quality control, the 16S rRNA V3-V4 hypervariable region was sequenced on the Illumina MiSeq platform system. The software system Quantitative Insights Into Microbial Ecology (QIIME2.2019.10) was used to conduct the quality filter of raw data. Microbial community analyses were conducted with R (version 4.1.1) vegan package as well as the functional prediction by PICRUSt2 by referenced scripts (60). The detailed protocol is shown in Text S1.

### Statistical analysis

Statistical significance was assessed using the Student’s t-test, one-way analysis of variance (ANOVA) followed by Tukey HSD multiple comparisons. All the data were confirmed to fit into Gaussian distribution by Shapiro–Wilk test for normality, and analyses were performed using Prism8 software (GraphPad Software, La Jolla, CA).

### Data availability

Sequencing data have been submitted to NCBI Sequence Read Archive (SRA) under accession number <u>SRP352833</u>.

## ACKNOWLEDGMENTS

This research was funded by the Ministry of Science and Technology, R.O.C. under grant number 110WFA0810232. We thank Dr. Po-Yu Liu for providing the custom R scripts for data analyses and Dr. Yu-Wei Wu for critically reviewing the manuscript.

## SUPPLEMENTAL MATERIAL

**FIG S1.**
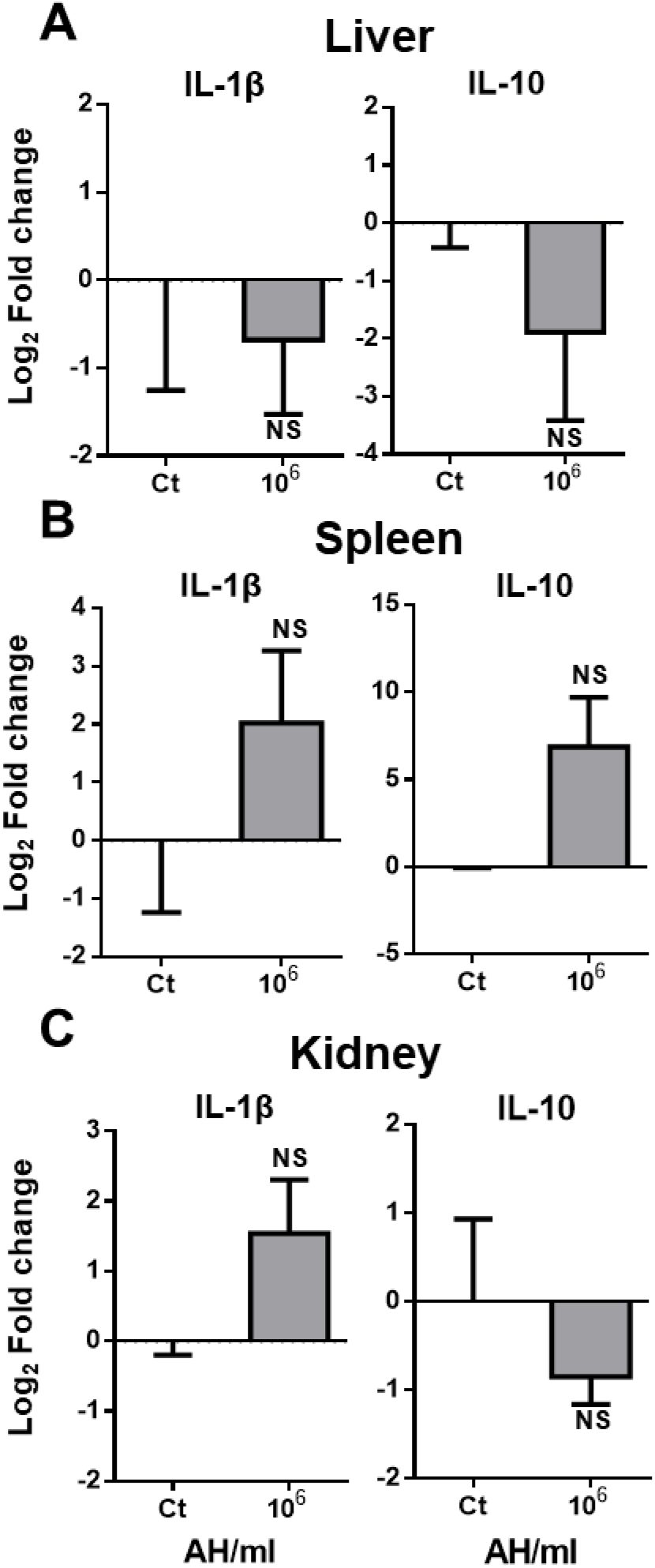
The immune response of liver, kidney, and spleen to AH challenge. The expression level of IL-1β and IL-10 in liver, spleen, and kidney was examined by qPCR. (A) Liver, (B) Spleen, and (C) Kidney in 10^6^ AH-challenged fish were measured compared to non-challenged control fish. N=3 fish in each group. Statistical significance was determined by Student’s t-test. (***p ≤0.001; **p≤0.01; *p≤0.05)

**FIG S2.**
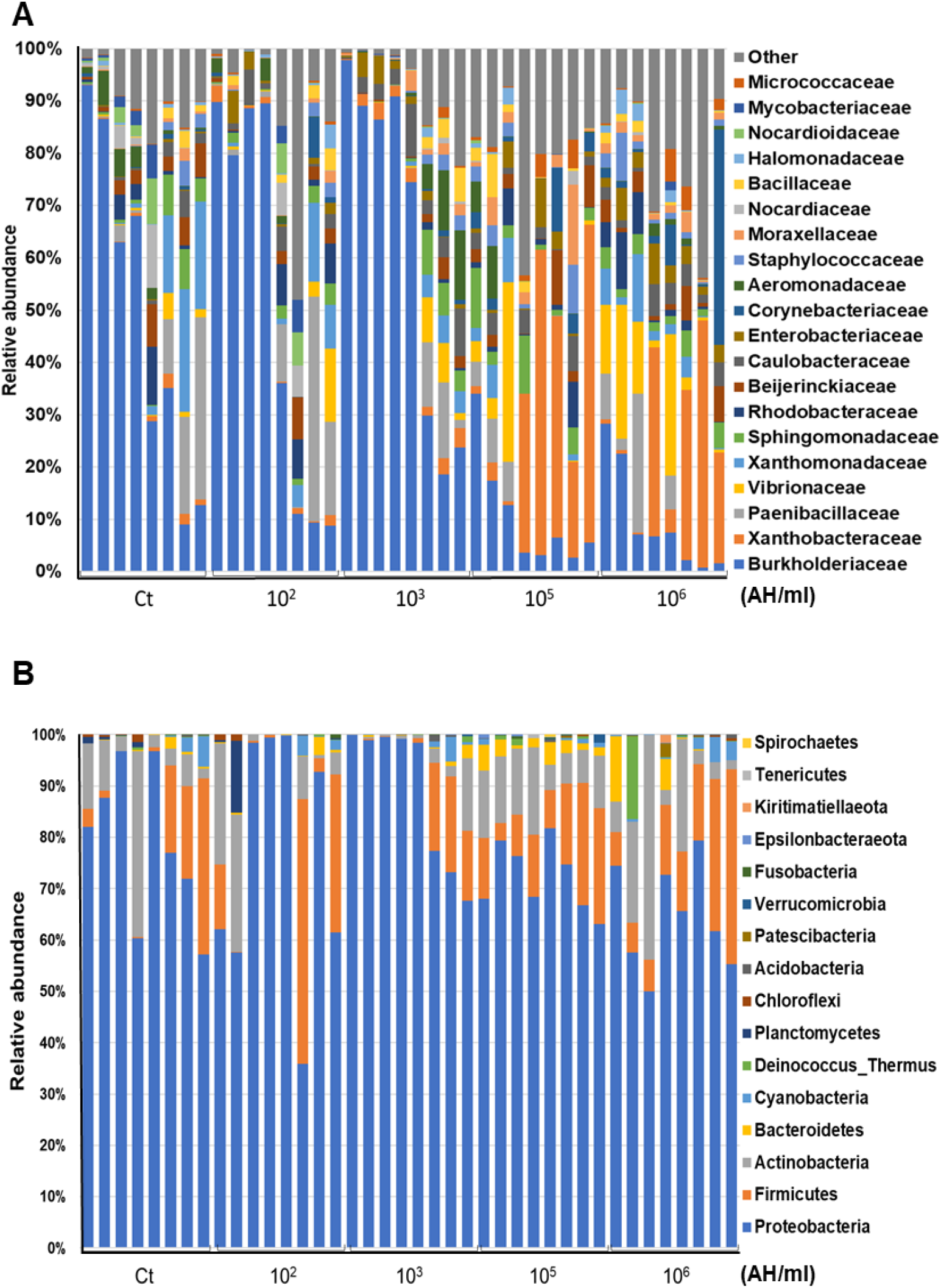
The skin mucosal microbiota composition of AH*-*challenged fish. The skin mucus of AH-challenged and non-challenged fish was scraped, and the microbial DNA was extracted, followed by 16S rRNA gene sequencing. The relative abundance of the skin mucosal microbiota composition for individuals in each group was shown as percentage at (A) phylum level (B) family level. N=8 fish in each group.

**FIG S3.**
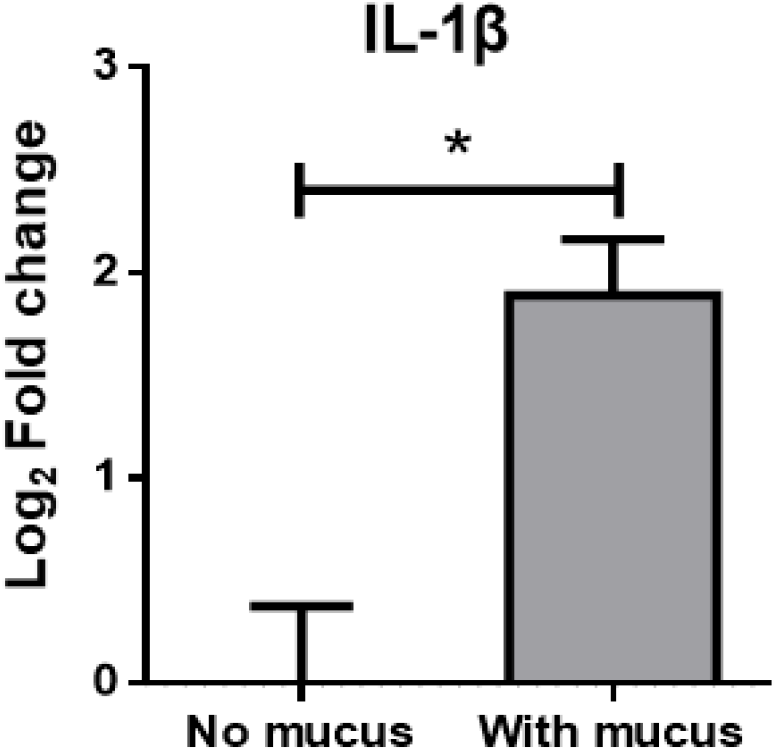
Striped catfish skin immune response to mucus. The fresh mucus was scraped and transplanted to the *ex vivo* skin model. After a 6-hr challenge, the fish skin was collected, and qPCR was performed on IL-1β expression level. Statistical significance was determined by Student’s t-test. (*, P ≤ 0.05; **, P ≤ 0.01; ***, P ≤ 0.001)

**FIG S4.**
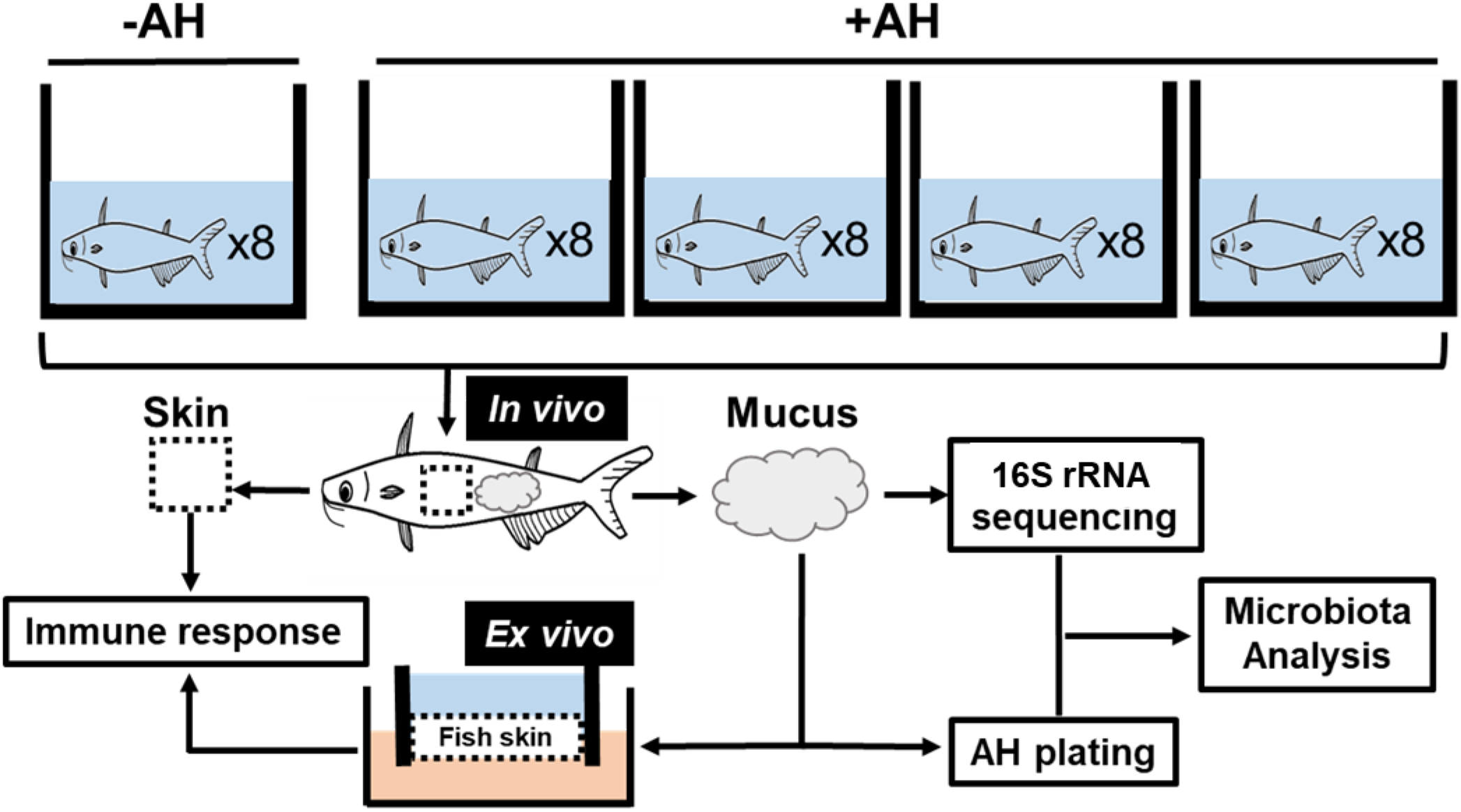
Experiment design and frameworks. Striped catfish were challenged with different concentrations of AH. After the challenge, the skin was removed and examined for immune response by qPCR. The mucus was used in microbiota examination by 16S rRNA gene sequencing analysis, Viable AH count by AH plating, and re-confirming the skin immune response by mucus transplantation into the *ex vivo* skin model system.

**FIG S5.**
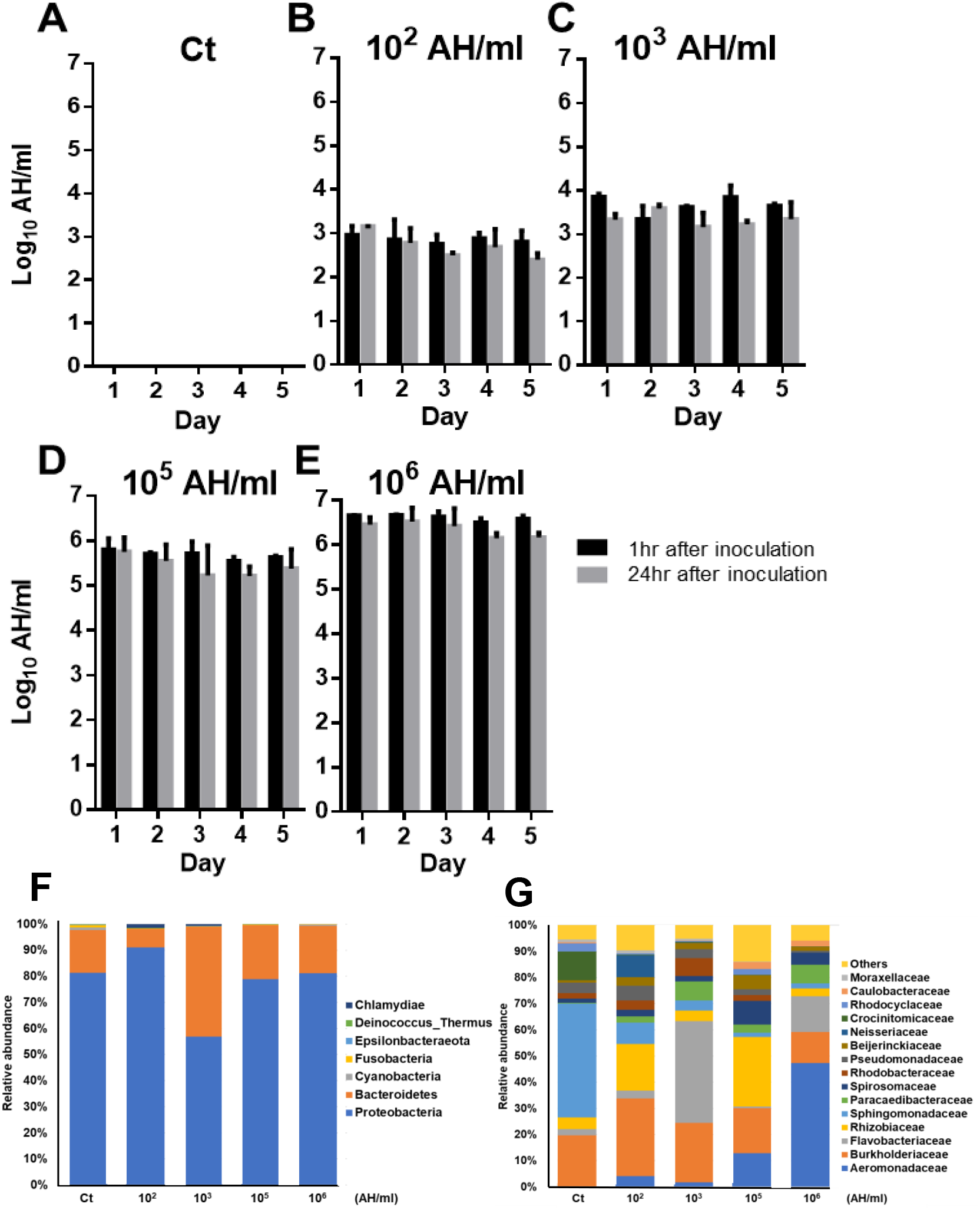
AH concentration in the five-day bath challenge period and the microbial composition of tank water at day5. AH concentration in water was measured by tank water plating during the five-day experiment period at two time points:1 and 24 hr after inoculation. At day5, the microbial DNA of tank water was extracted, followed by 16S rRNA gene sequencing. Shown are AH concentration of (A) Control, (B) 10^2^, (C) 10^3^, (D) 10^5^, and (E) 10^6^ AH/ml tanks and the relative abundance of the microbiota composition for experimental tank water at day5 was shown in each group as percentage at (F) phylum level (G) family level. Statistical significance was determined by One-way ANOVA followed by Tukey’s post-hoc test. (Different letters indicate significant differences at p < 0.05)

**TABLE S1.** AH was examined in liver, spleen and posterior kidney in both control and 10^6^ AH/ml challenge group.

| Group/Fish number | Liver |  |  |  |  | Spleen |  |  |  |  | Kidney |  |  |  |  |
| --- | --- | --- | --- | --- | --- | --- | --- | --- | --- | --- | --- | --- | --- | --- | --- |
|  | 1 | 2 | 3 | 4 | 5 | 1 | 2 | 3 | 4 | 5 | 1 | 2 | 3 | 4 | 5 |
| Ct | - | - | - | - | - | - | - | - | - | - | - | - | - | - | - |
| 10<sup>6</sup>AH/ml | - | - | - | - | - | - | - | - | - | - | - | - | - | - | - |

**TABLE S2.**
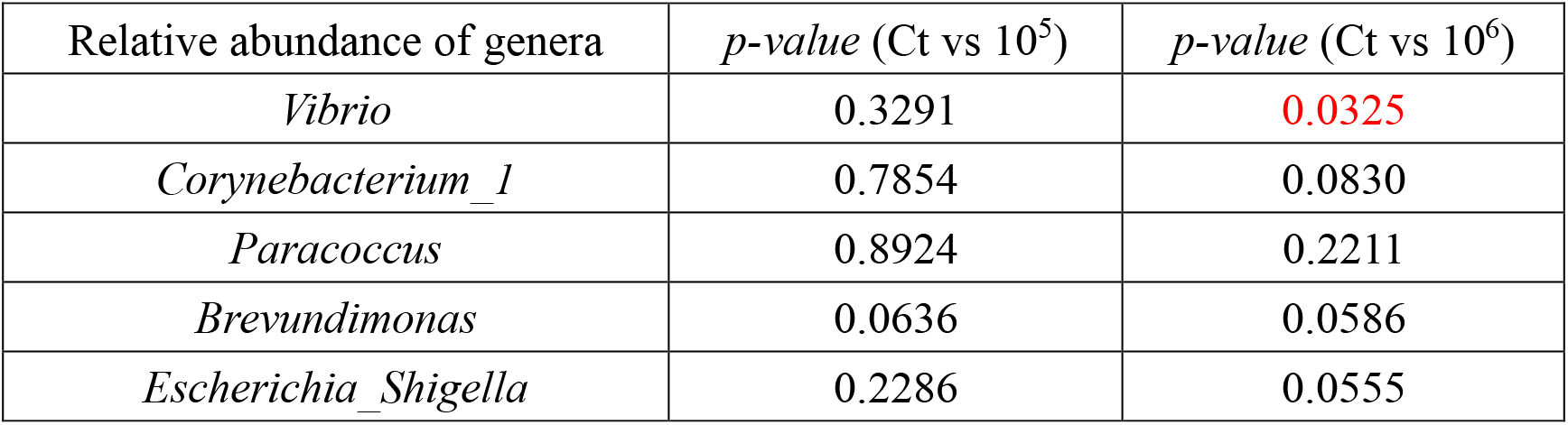
Statistical analysis was applied to determine the significant difference of relative abundance of bacteria genera between 10^6^ AH/ml to non-AH control and 10^5^ AH/ml group to non-AH control. (Student’s t-test, ***p ≤0.001; **p≤0.01; *p≤0.05)

**TABLE S3.** Anti-AH property of the skin mucus was examined by different treatments, included the positive control of tetracycline, the negative control of double-distilled water and the different concentration of mucus.

| *Aeromonas hydrophila*<br>concentration | Treatment |  |  |  |  |
| --- | --- | --- | --- | --- | --- |
|  | tetracycline<br>(50mg/μl) | double-distilled<br>water | mucus<br>(1X) | mucus<br>(10X) | mucus<br>(1X/100) |
| 10<sup>5</sup>AH/ml | + | - | - | - | - |
| 10<sup>6</sup>AH/ml | + | - | - | - | - |
| 10<sup>7</sup>AH/ml | + | - | - | - | - |

**TABLE S4.**
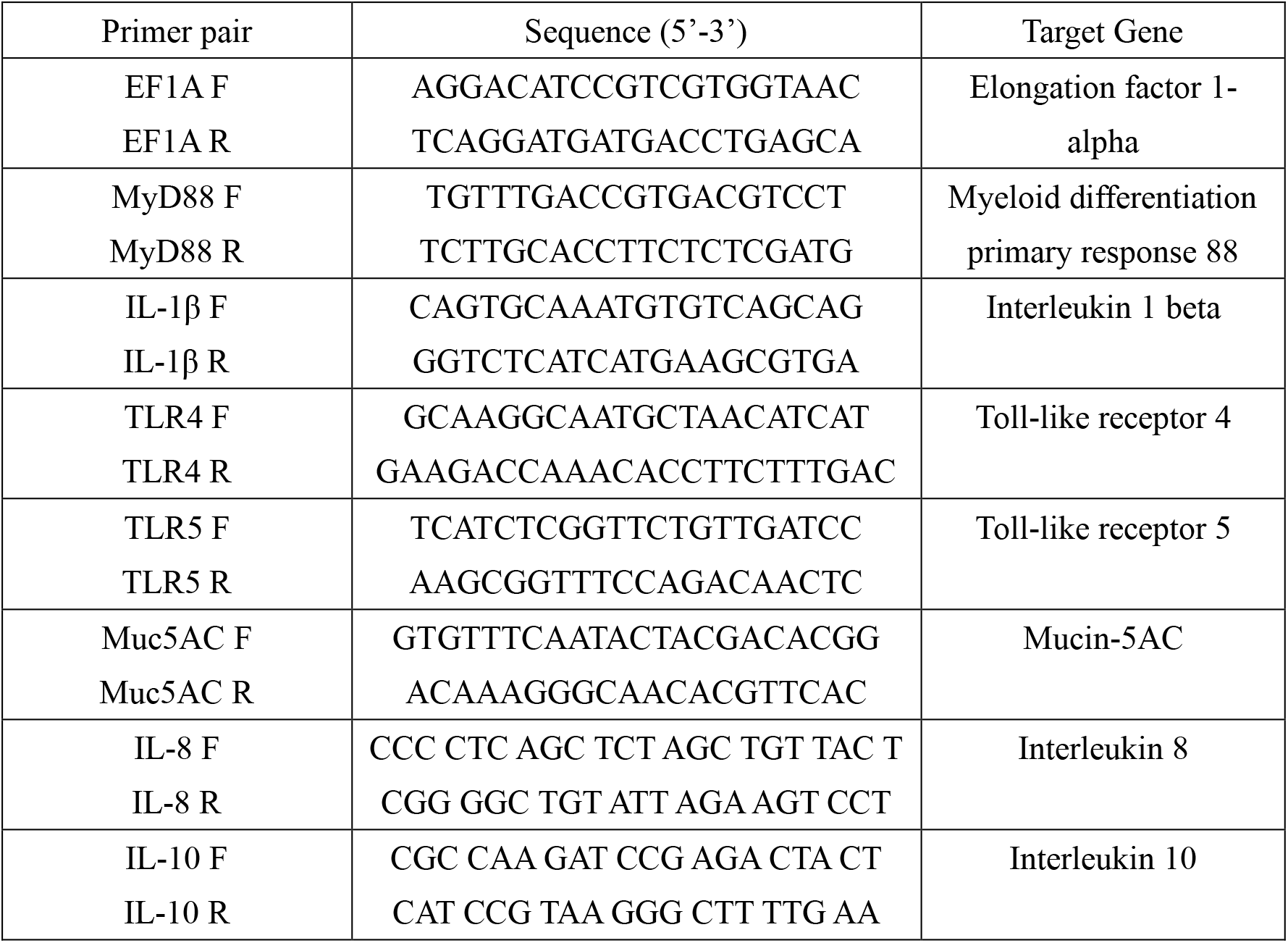
Primer pairs used in this research TEXT S1 Supplemental Materials and Methods.

### TEXT S1 Supplemental Materials and Methods

#### Total RNA extraction and cDNA preparation

Each total RNA sample from the skin, liver, spleen, and kidney of striped catfish were extracted with the TriPure Isolation Reagent (Roche, Mannheim, Germany) following the manufacturer’s instructions. Extracted RNA samples were dissolved in 30 μl of sterilized DDW and stored at −80 °C after isolation. The concentration and quality of extracted RNA were measured by DS-11 Series Spectrophotometer (DeNovix, USA). Purified RNA with an A260/A280 ratio between 1.8 and 2.0 was used for all RNA experiments. First-strand cDNA was synthesized from 2 mg of total RNA by M-MLV Reverse transcriptase (Promega, USA) following the manufacturer’s instructions. The successful construction of cDNA library was determined by 1.5% agarose gels containing the Safeview DNA stain (GeneMark, Taiwan)

#### Real-Time PCR (qPCR)

Primer pairs were designed using Universal Probe library website and shown in Table S4. All the cDNA products were utilized for the qPCR reaction using the GoTaq qPCR Master Mix on a CFX96 real-time PCR Detection System (Bio-Rad, CA). The thermal cycling profile consisted of an initial denaturation at 95°C (for 2 min), followed by 40 cycles of denaturation at 95°C (3 sec), an appropriate annealing/extension temperature (60°C, 30 sec). An additional temperature ramping step was utilized to produce melting curves of the reaction from 65°C to 95°C. The housekeeping gene EF1-α was set as the reference gene, relative fold changes were calculated based on the cycle threshold (Ct) values generated by qPCR. Expression differences between control and treatment groups were assessed for statistical significance using a randomization test in the GraphPad Prism. The numbers in y-axes correspond to the Log_2_ fold increase compared to the non-AH challenge group.

#### Skin mucus microbial DNA extraction

*P. hypophthalmus* skin mucus about 20 μl was homogenized in 100 μl of TE buffer (1 M, pH 8 Tris-Cl; 0.5M, pH 8 EDTA) and 100 μl lysozyme (50 mg/ml) then incubated at 37°C for 30 min. Then, TE buffer, 10% SDS, and proteinase K (20 μg/ml) were added, and the sample was incubated at 56°C for 1 hr. After that, 5 M NaCl and phenol:chloroform:isoamyl alcohol (25:24:1) was added to the sample and centrifuged. chloroform:isoamyl (24:1) was added to the collected supernatant and was also centrifuged. The DNA was precipitated with 90% ethanol, put in -20°C overnight, and centrifuged the next day again. The DNA pellet was rinsed with 70% ethanol and centrifuged once more, air-dried in a laminar flow hood, and resuspended in TE buffer or double-distilled water. All centrifugation steps were performed at 14000 rpm, 5 min, and 4°C.

#### 16S rRNA gene sequencing and community analyses

Skin mucus microbial DNA was extracted and assessed photometrically using DS-11 Series Spectrophotometer (DeNovix, USA). The V3-V4 hypervariable region of 16S rRNA genes was amplified by the primer set 341F (5′-CCTACGGGNGGCWGCAG-3′) and 805R (5′-GACTACHVGGGTATCTAATCC-3′). After preparation and quality control, the 16S rRNA libraries were sequenced on the Illumina MiSeq platform system. After sequencing, we obtained 50000-100000 raw reads from the original samples. Silva classifier was applied for the taxonomic assignment. The software system Quantitative Insights Into Microbial Ecology (QIIME2.2019.10) was used to conduct the quality filter of raw data. Then these sequences were clustered into operational taxonomic unit (OTU) picking that can apply to taxonomy assignment, the computation of α and β diversity measures, and other analyses. Microbial community analyses were conducted with R (R version 4.1.1) package vegan. A one-way ANOVA test in R software44, with α=0.05, was used for all statistical analyses and Tukey HSD test for post-hoc comparisons. To normalize sequencing output among samples, we rarefied the ASV/OTU table around 1000 reads per sample. Alpha diversity was analyzed based on the OTU table, which was present in 50% of the sample in each group, Shannon index was calculated by ‘ diversity ’ function, species richness (S) was counted by ‘specnumber’function and species evenness (J) was calculated by following formula J=H’/lnS’. For beta diversity, dissimilarities among microbial communities were measured by Bray-Curtis distance and conducted with principal coordinates analysis (PCoA) based on the bacterial family classification level. LEfSe was applied for identifying featured mucus microbes among striped catfish by using relative abundances of OTUs. The featured OTUs were first tested and detected with a Kruskal-Wallis test (p-value cutoff=0.1); then, linear discriminant analysis (LDA) was conducted for selecting featured OTUs by effect size (absolute value of the logarithmic LDA score=2). The selected featured OTUs which belong to each body-size class were visualized with a cladogram, based on their taxonomy.

#### Functions prediction of microbial community

PICRUSt2 was used to predict the metagenome to evaluate potential functions of striped catfish’ skin microbiota. Copy number adjusted OTU tables were used for predicting Kyoto Encyclopedia of Genes and Genomes (KEGG) Orthology (KO) abundances. The KOs were categorized by KEGG pathway database and specific metabolic module/reactions of interest (i.e. cellulose degradation module in starch and sucrose metabolism pathway). Pathway/module topology analysis was conducted with the R (R version 4.1.1) package path view.

#### *Ex vivo* fish skin model system

A striped catfish model system was applied in this research. The striped catfish skin tissue was removed from the fish by scalpel and immediately immersed into cold L-15 medium (SIGMA, U.S.A.) supplemented with 10% fetal bovine serum, 2% gentamycin solution (SIGMA, U.S.A), 1X antibiotic-antimycotic (Biowest, U.S.A.). After cutting into squares of approximately 10×10 mm, skin tissue was fixed in the upper plastic crown using a fine rubber band and mounted with the lower plastic crown. The tissue and the crown were gently submerged in the culture medium in a 24-well culture plate. The plate was cultured in a CO_2_-free incubator at 25°C. The culture medium was changed every two days until further experiments were conducted. The media was replaced with non-antibiotic media 24 hr before the infection experiment.

